# Vagus nerve mediated liver-brain axis is a major regulator of the metabolic landscape in the liver

**DOI:** 10.1101/2023.11.16.567412

**Authors:** Camila Fatima Brito, Roberta Cristelli Fonseca, Lucas Rodrigues-Ribeiro, Bruna Fernandes Vaz, Gabriel Sousa Silva Tofani, Ariane Barros Diniz, Paola Fernandes, Núbia Alexandre Melo Nunes, Rafaela Miranda Pessoa, Amanda Carla Clemente Oliveira, Valbert Nascimento Cardoso, Simone Odília Antunes Fernandes, Maristela Poletini Oliveira, Jacqueline Isaura Alvarez-Leite, Gustavo Batista Menezes, Adaliene Versiani M Ferreira, Vladimir Gorshkov, Frank Kjeldsen, Thiago Verano-Braga, André Gustavo Oliveira

## Abstract

**Background:** The liver serves as a major energetic reservoir for other tissues and its metabolic function is controlled by humoral and neural factors. The vagus nerve innervating the gastrointestinal tract plays an important role in regulating peripheral metabolism and energy expenditure. Although the liver receives vagus nerve fibers, the impact of this circuitry in the regulation of hepatic metabolism is still poorly understood.

**Methods:** Herein, we used a combination of quantitative proteomics and *in vivo* imaging techniques to investigate the impact of the vagus nerve on liver metabolism.

**Results:** Vagus nerve shapes the metabolic framework of the liver, as surgical ablation (vagotomy; VNX) of this circuitry led to a significant alteration of the hepatic proteome landscape. Differential protein expression and pathway enrichment analyses showed that glycolytic and fatty acid biosynthesis were increased following VNX, whereas β-oxidation was decreased. This metabolic shift facilitated lipid accumulation in hepatocytes. Furthermore, VNX worsened liver steatosis following high-carbohydrate or high-fat dietary challenges.

**Conclusions:** This study describes the liver-brain axis mediated by the vagus nerve as an important regulator of the hepatic metabolic landscape.

**Highlights:**

- Vagus nerve is a novel regulator of the hepatic metabolic landscape.
- Ablation of vagus nerve neural circuit by vagotomy resulted in a metabolic shift towards glycolysis and fatty acid biosynthesis.
- Lipid accumulation was increased in vagotomized mice fed with a standard diet.
- Liver steatosis was increased following dietary challenges with high-carbohydrate or high-fat diets.
- Vagus nerve can be a promising new target for NAFLD treatment.

## 1. Introduction

Heterotrophic organisms depend on nutrients to fuel metabolism and support life [1]. Following digestion and absorption in the intestines, nutrients and microbiota-derived metabolites enter the liver, a central metabolic hub in the organism, through the hepatic portal circulation.

The hepatic parenchyma is a highly specialized microenvironment that supports and integrates several key metabolic frameworks and reactions [2] that are important to produce and store the energy used to sustain physiological functions at the cellular, tissue and organ levels [3]. Liver metabolism is highly dynamic and is regulated by nutrient availability and humoral factors, which confer the flexibility needed for adaptation to fed and fasting states. In addition, a growing body of evidence suggests that the autonomous central nervous system also plays an important role in coordinating hepatic metabolic functions [4].

The vagus nerve is an important source of autonomic innervation to the gastrointestinal tract [5]. Because it contains both afferent and efferent fibers, the vagus nerve conveys a bidirectional circuitry bridging these organs and the brain. Notably, a great number of vagal fibers derived from the hepatic branch innervate the liver and thus may couple nutrient ingestion with metabolism. The concept is that vagal afferent innervation senses mechanical distention of the digestive tube following food ingestion, paralleled by a vagal-mediated neuroepithelial circuit that senses nutrients within the intestinal lumen [6,7]. In addition, a role for hepatic vagal innervation in nutrient sensing has been proposed, as portal infusion of glucose, amino acids and lipids are shown to increase hepatic vagal activity [8–11]. These stimuli are integrated in different brain areas and are classically involved in learning of food preferences, controlling meal size (satiation), as well as proper digestion and absorption of nutrients [12,13]. Not surprisingly, the disruption in the neurophysiology of gastrointestinal-brain axis has been associated with metabolic diseases. In this regard, the reduction of vagus nerve sensitivity or plasticity resulted in hyperphagia and led to obesity [14]. In line with these findings, electrical stimulation of the vagus nerve reduced food intake, body weight/fat and increased energy expenditure [15–17]

Here, we show that in addition to sensing mechanical and nutrient stimuli, the vagus nerve displays an important role in shaping the metabolic framework of the liver. Furthermore, we show that the disruption in this circuitry shifts hepatic metabolism and facilitates lipid accumulation in hepatocytes.

## 2. Materials and Methods

### 2.1. Ethical approval

All experimental procedures were performed in accordance with protocols approved by the institutional Animal Use Committee and in compliance with the Brazilian legislation of animal care and experimentation.

### 2.2. Mouse models and surgical procedure

Male C57/Bl6 mice (8-10 weeks old) used in this study were housed under a 12-hour light/dark cycle at 25 °C, with unrestricted access to food and water. All the experiments were performed mid through the light phase at ZT6 ± 30min (ZT; *Zeigtgeber Time*) as the circadian light cycle influences the majority of metabolic pathways in the murine liver [18].

Vagotomy (VNX) was performed in mice anesthetized with ketamine/xylazine (i.p.; 80mg/Kg and 15mg/Kg, respectively). After a midline incision in the cervical region, the left cervical trunk of the vagus nerve was exposed, ligated with a 4-0 silk and sectioned [19]. Sham mice were submitted to similar procedures without nerve transection. All animals were allowed to recover for 7 days before other experiments.

### 2.3. Proteomics

To get an overview of the proteins expressed in our phenotype, we performed the proteomics assay, which also served as the basis for understanding possible subsequent physiological changes in our model. After seven days of recovery from surgery, we collected the livers. Samples were washed with physiological saline, frozen in liquid nitrogen and later stored in a freezer at −80° C.

#### 2.3.1. Liver tissue preparation

Tissues were snap-frozen in liquid nitrogen and grounded to powder in a mortar chilled with liquid nitrogen using a pestle. Tissues were stored at −80 °C until further use. About 6 milligrams (mg) of sample was resuspended in 100 µL of lysis buffer containing 6M Urea, 2M Thiourea, 40mM 2-chlorocetamide, 20mM TEAB, 100mM TCEP and protease and phosphatase inhibitors. The samples were sonicated at 40Hz, 3x for 15 seconds in ice-cold for DNA fragmentation. Then, the samples were incubated at 28°C and 650 rpm for 2h for proper protein lysis, reduction and alkylation. After that, protein quantification was quantified using Qubit assay (Thermo Fisher Scientific, USA) and 250μg of proteins of each sample were collected for protein digestion. Prior digestion, samples were diluted with TEAB 50mM to reach 0.6M urea and pH 8. Trypsin was added at 1:50 (w/w) (enzyme-to-protein ratio) and protein digestion was carried out at 25 °C for 16 h. Enzymatic reaction was quenched with TFA 1% (v/v) (final concentration).

Samples were desalted using self-made micro-columns packed with OLIGO R2 and R3 Reversed-Phase Resin (Thermo Fisher Scientific, USA). Desalted peptides were dried in a speedvac and stored at −20 °C until further use.

#### 2.3.2. Peptide isotopic labeling

Peptides were labeled according to the “On-column” dimethyl labeling protocol [20], with minor modifications. Briefly, the peptides were reconstituted in 1mL of 5% (v/v) formic acid (CH_2_O_2_). SepPak columns (Waters, USA) were: (i) activated with 4 mL of 100% ACN; (ii) conditioned with 4 mL of solution A (0.6% acetic acid (v/v)); (iii) loaded with the samples; (iv) washed with 4 mL of solution A; (v) incubated with the respective labeling reagent for 30 minutes. Sham and VNX groups were labeled with different labeling reagents (Sham = light and VNX = medium). The isotopic labeling reagents contained 500 µL of 50 mM sodium-phosphate buffer (pH 7.5), 250µL of 0.6M sodium cyanoborohydride (NaBH_3_CN) and 250µL of a 4% (v/v) solution containing the respective formaldehyde: CH_2_O (light) or CD_2_O (medium); (vi) washed with 4 mL of solution A; (vii) eluted with an increasing gradient (20%, 50%, 80% and 100% (v/v) ACN + 0.6% (v/v) acetic acid). Samples were dried in a speedvac and reconstituted in 0.1% TFA (v/v) and quantified using Qubit (Thermo Fisher Scientific, USA). The labeling efficiency was checked using the MALDI-TOF mass spectrometer (Autoflex III, Bruker) with the m/z range of 780-2500 Da. Then, the samples from the light and medium groups were combined in the same tube with the same ratio of 1:1 labeled peptides (Sham:VNX). Finally, samples were dried using a speedvac and stored at −20°C.

#### 2.3.3. Mass spectrometry analysis

Fractions were resuspended in 0.1% (v/v) formic acid (solvent A) and the peptides were separated and analyzed using an EASY-nLC system (Thermo Fisher Scientific, USA) with a two-column system setup coupled to a Q-Exactive HF mass spectrometer (Thermo Fisher Scientific, USA). The pre-column was 3 cm length x 100 µm i.d. and was packed with 5 μm particles (Reprosil-Pur C18-AQ, Dr. Maisch GmbH, Germany). The analytical column was 18 cm length × 75 µm i.d. and was packed with 3 μm particles (Reprosil-Pur C18-AQ, Dr. Maisch GmbH, Germany). For the peptide separation, the following chromatographic gradient was used: 1–3% solvent B (95% (v/v) acetonitrile + 0.1% (v/v) formic acid) in 0-3 min; 3–28% solvent B in 45 min; 28–45% solvent B in 10 min; 45–95% solvent B in 3 min at 250 nL/minute. For the MS analysis, the instrument was operated in positive polarity and DDA mode. The eluted peptide ions were resolved (MS1 mass range = m/z 400–1400) at 120000 resolution (at m/z 200). The MS1 AGC target settings were set to allow accumulation of up to 3 × 10^6^ charges for up to 120 ms injection time. The 20 most intense peptide ions (Top 20) were selected with an isolation window of 1.2 Th and the normalized collision energy for HCD fragmentation was set to 32. For MS2 AGC target settings were set to allow accumulation of up to 10^5^ ions for up to 120 ms. The fragment ions were resolved with 15000 resolution (at m/z 200) and selected precursor ions were excluded. Spectra files (.raw) were viewed in Xcalibur v3.0 (Thermo Fisher Scientific, USA).

### 2.4 Proteomics data analysis

The raw spectra (.raw) were analyzed with MaxQuant 1.5.8.3. Spectra were searched against the SwissProt *Mus musculus* FASTA database (downloaded October 2017) using the Andromeda search engine [21]. Search parameters included a 20 ppm tolerance for the first search and a 4.5 ppm tolerance for the main search. Dimethyl labeling was set using the following modifications: “Dimethyl-Lys0” and “Dimethyl-Nterm0” for light labeling, and “Dimethyl-Lys4” and “Dimethyl-Nterm4” for medium labeling. Trypsin was chosen as the enzyme with a max of two missed cleavage sites. Carbamidomethyl of cystein was set as a fixed modification, and oxidation of methionine and acetylation of protein N-terminus were set as variable modifications. “Match between runs” was enabled with a matching time window of 0.7 min and an alignment time window of 20 min. Peptide identification required at least 1 unique peptide and was filtered to 0.01 false discovery rate (FDR).

Protein intensity was log2-transformed and then normalized by the column median and row mean. Reverse and contaminant proteins were removed using Perseus software. Statistical analysis one-way ANOVA (p-value < 0.05) was applied to find differentially regulated features with FDR 0.05 strategy to adjust the p-values using DanteR. To explore the data within the Kegg database context, we used Functre2 [22]. Kegg functional enrichment and further data visualization were performed in R (version 4.1.3) and the following R-packages: ClusterProfiler [23], EnhancedVolcano and pheatmap. Protein network analysis was performed using String and ClueGo [24,25] for Cytoscape (version 3.7.2).

### 2.5. Locomotor activity

A telemetry sensor was surgically placed in the abdominal cavity of VNX and control animals. The locomotor activity was telemetrically monitored through the signal emitted by the capsule implanted in the abdominal cavity and received by the signal receiver under the experimental vats connected to an acquisition software. Data was acquired in continuous mode and subsequently processed. Telemetry records were made for 14 days.

### 2.6. Intestinal Motility

VNX or Sham animals (n = 5 per group) received, by gavage, 300μL of activated carbon solution (10% v/v of activated carbon:5% v/v of gum arabic), at ZT6 ± 30min. Motility was determined in isolated intestines 20 minutes after gavage by measuring the distance traveled by the activated carbon solution and dividing it by the value of the total length of the intestine. The result was expressed as a percentage of the total length of the small intestine.

### 2.7. Intestinal Absorption

All animals received by gavage 100μL of solution containing diethyleneaminopentacetic acid labeled with 18.5 MBq 99mTechnetium (99mTc-DTPA). Four hours after gavage, blood samples were collected and submitted to radiation determination (cpm) in a gamma radiation counter (Wallac Wizard Gamma counter, PerkinElmer, USA). The results obtained were compared with the standard of 99mTc-DTPA and calculated as the percentage product of blood cpm times 100 cpm of administered dose.

### 2.8. Dietary challenges

To analyze the temporal evolution of diet-induced metabolic changes, we introduced a high-refined carbohydrate (HC) or high-fat (HFD) diet in vagotomized animals and their control. The macronutrient composition of the HC diet (4.4 kcal/g) was 74.2% carbohydrate, 5.8% fat and 20% protein. It is important to note that the HC diet contains at least 30% refined sugars, mainly sucrose. Standard diet (4.0kcal/g) composition was 65.8% carbohydrate, 3.1% fat and 31.1% protein (Nuvilab, Quimtia, Brazil). The animals were fed standard laboratory chow or experimental diet for a period of 8 weeks after vagotomy.

In the HFD diet model, mice were fed standard laboratory chow or high-fat chow for a period of 4 weeks after vagotomy. High-fat diet (7.0 kcal/g) was composed of 24.5% carbohydrate, 61.1% fat and 14.5% protein. At the end of dietary treatment, animals were euthanized. Liver samples were frozen in liquid nitrogen and stored at −80° C until processing.

### 2.9. Intravital microscopy

The intravital confocal microscopy images were performed as described [19,26]. In summary, mice from the Sham and VNX groups were anesthetized with an intraperitoneal injection of ketamine (80 mg/kg) and xylazine (15 mg/kg), at the time of ZT6. A laparotomy was performed to expose the liver and then the left lobe was collected and placed in a petri dish containing saline. To visualize the lipid droplets in vivo, the Bodipy dye (1.5 μg/mouse diluted in DMSO, Thermo Fisher Scientific, USA) was placed directly in the left lobe of the liver collected after surgery. Images were obtained using a Nikon A1R confocal microscope (Nikon Instruments, USA). Images were acquired with a 20x Plan Apo objective and analyzed using the ImageJ software.

### 2.10. Statistical analysis

Differences between two samples were analyzed for significance using unpaired two-tailed Student’s t-test. Differences between three or more groups were analyzed using One-way ANOVA with Newman-Keuls Multiple Comparison Test. Values are from multiple biological replicates within an experiment and reported as the mean±standard deviation and plots that highlight the distribution of individual data. Statistical significance was set at p<0.05.

## 3. Results

### 3.1. Vagotomy leads to hepatic proteome remodeling

To elucidate the processes regulated by the vagus nerve in the liver, we first performed quantitative proteomics that allowed a comparison of protein relative expressions between vagotomized mice (VNX) and sham-operated control animals (Sham). In total, our proteomic analysis of five biological samples for each group allowed the identification and quantification of 2121 proteins (Supplementary Table S1). Importantly, aiming to increase robustness of downstream analysis, we did not exclude any biological replicate. Furthermore, if a certain protein was not identified in all samples, it was not included in our study.

Principal component analysis (PCA) showed that Sham and VNX animals had distinctly different proteomes, as both groups clustered apart (Fig.1A). We also identified 453 differentially expressed proteins between VNX and Sham groups, corresponding to approximately 21% of the total proteins, of which 235 were downregulated and 219 upregulated following vagotomy (Fig.1B-C). Taken together, our results indicate that the proteome of the liver was remodeled following vagotomy.

**Figure 1:**
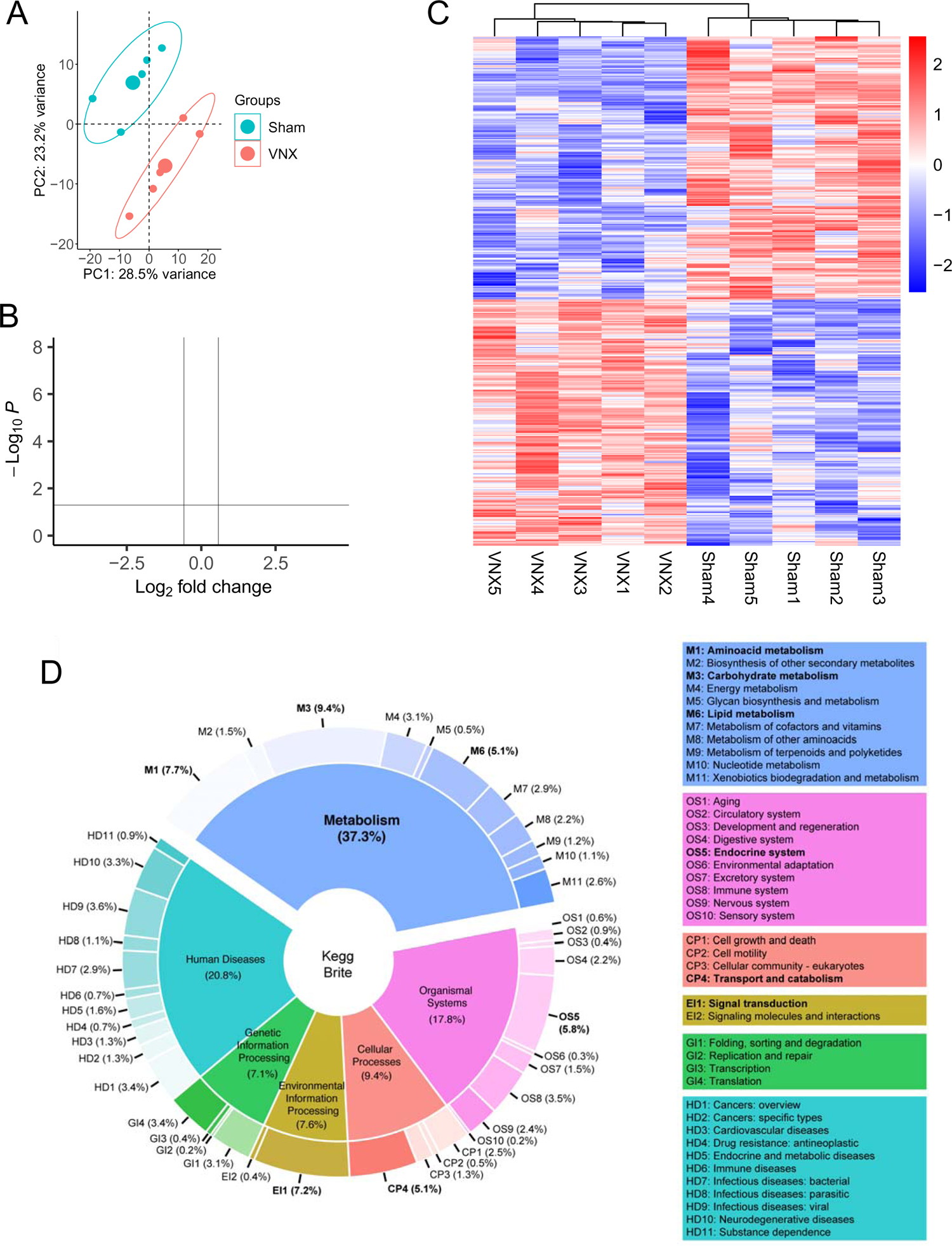
Liver proteome remodeling following vagotomy. **(A)** PCA of relative protein expression in VNX and Sham animals. **(B)** Volcano plot of protein expression. Blue and red colors respectively represent statistically decreased and increased expression in VNX mice compared to Sham animals (n=5/group; p<0.05, FDR adjusted-p<0.05). **(C)** Significantly regulated protein patterns in individual samples. **(D)** Summary of significantly regulated proteins mapped onto the KEGG database. VNX = vagotomy.

### 3.2. Metabolism is a major target of the vagal neural circuit in the liver

To gain insights into the biological functions associated with the differentially abundant proteins following VNX, we performed an exploratory analysis using the hierarchical context of the KEGG Brite database. In the first Brite level, these proteins covered 37.3% of biological objects classified within the “Metabolism” category, followed by objects categorized into “Human Diseases” (20.8%), “Organismal Systems” (17.8%), “Cellular Processes” (9.4%), “Environmental Information Processing” (7.6%) and “Genetic Information Processing” (7.1%) (Fig.1D). In a further classification level, we observed that the different abundant proteins covered functions classified into key metabolic and/or regulatory processes, including “Carbohydrate metabolism” (9.4%), “Amino acid metabolism” (7.7%), “Lipid metabolism” (5.1%), “Energy metabolism” (3.1%), Endocrine system (5.8%) and “Signal transduction” (7.2%) (Fig.1D).

Next, we mapped the proteins onto a hierarchical dendrogram that comprehends all known biological functions of the KEGG database, where each layer of nodes is mapped to a level of hierarchy. In addition, we also mapped the sum of the inverse p-value to the radius of each node, so that the node size could facilitate the visualization of putative functional clusters altered by vagotomy. Based on our previous results, we set the root of the dendrogram in Metabolism, as it was the Brite category with more coverage. We confirmed that carbohydrate, lipid and amino acid metabolism were the main metabolic processes impacted by vagotomy (Fig.2A). This analysis also allowed the investigation of biological pathways within these categories and revealed that core metabolic pathways and subpathways (or modules) in the liver depend on the proteins regulated by the vagus nerve. In this sense, we observed that glycolysis, including Embden-Meyherhof pathway (glycolysis => pyruvate and compounds); gluconeogenesis (oxaloacetate => fructose-6P); fatty acid biosynthesis (initiation and elongation) and fatty acid degradation (beta-oxidation) were important targets regulated by the vagus nerve and that VNX potentially disrupted the hepatic metabolic flux (Fig.2A).

**Figure 2:**
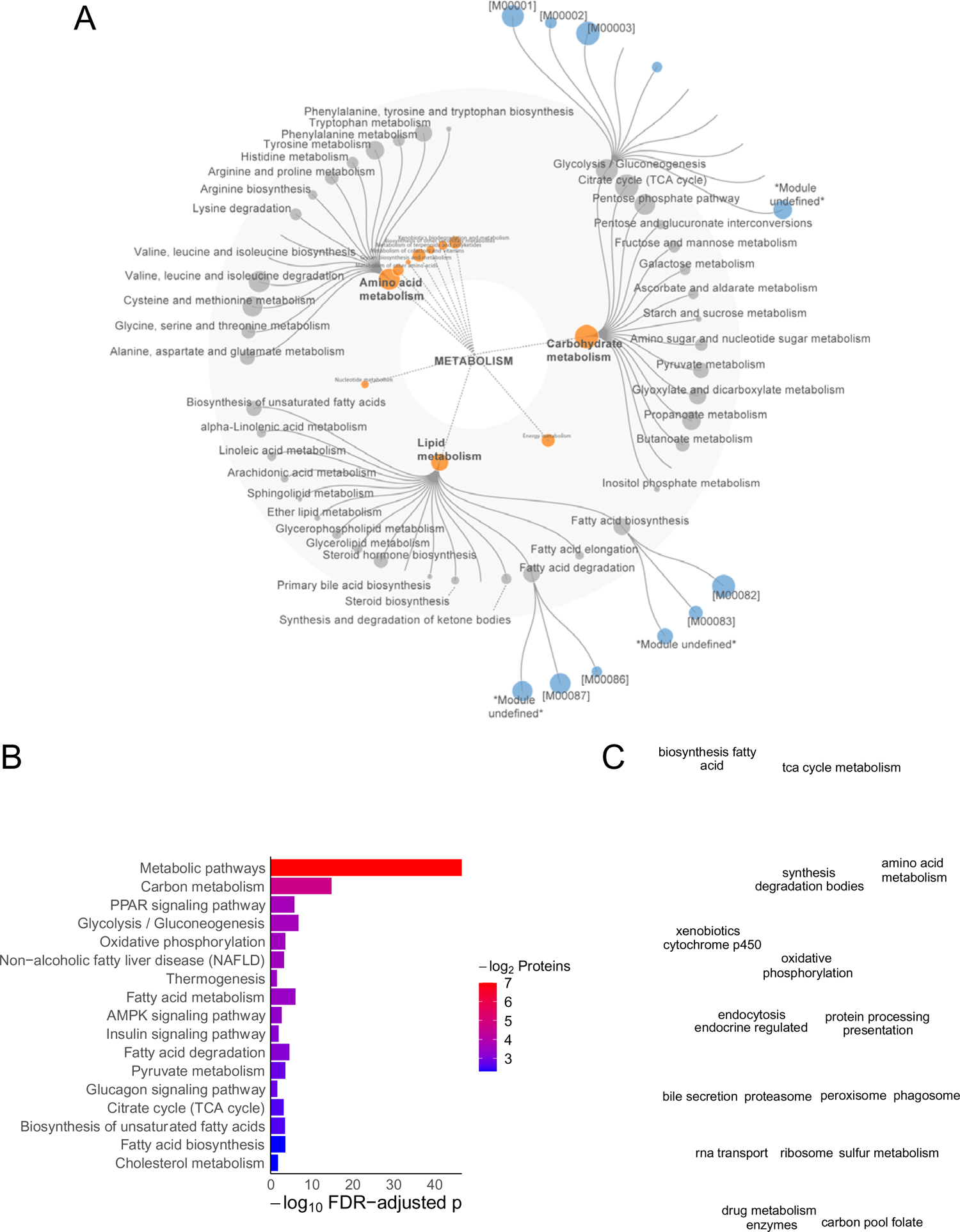
Functional enrichment analysis reveals metabolism as a major target of vagus nerve circuitry. **(A)** Functional mapping, **(B)** enrichment analysis and **(C)** pathway clustering network of differentially abundant proteins following vagotomy. Node size in (A) corresponds to the inverse FDR adjusted p-values. M00001 = Glycolysis (Embden-Meyerhof pathway), glucose => pyruvate; M00002 = Glycolysis, core module involving three-carbon compounds; M00003 = Gluconeogenesis, oxaloacetate => fructose-6P; M00082 = Fatty acid biosynthesis, initiation; M00083 = Fatty acid biosynthesis, elongation; M00086 = beta-Oxidation, acyl-CoA synthesis; M00087 = beta-Oxidation. In (B), enriched pathways were considered when FDR<0.05. Node in (C) represents enriched pathways and text corresponds to the word count of terms associated with the pathways within each cluster.

Not surprisingly, a functional enrichment analysis using the KEGG pathway database showed that the majority of the enriched pathways were indeed related to metabolic processes (Fig.2B). These results agree with our previous analyses. The network reconstruction of the enriched terms revealed that the pathways formed functional modules, as they clustered together and the word tag clouds of their annotations highlighted three major modules associated with glycolysis/gluconeogenesis, lipid metabolism and TCA cycle (Fig.2C). Taken together, our results suggest that the vagus nerve is an important player in the control of liver metabolism.

### 3.3. Vagotomy results in a metabolic shift towards fatty acid biosynthesis in the liver

We then reconstructed the metabolic framework of proteins within the pathways comprising the most represented functional modules: glycolysis/gluconeogenesis and lipid metabolism. The protein networks revealed that the vagus nerve has a broad regulatory effect, as it regulates the expression of multiple enzymes in each pathway, thus imbalancing the global hepatic metabolic interactome (Fig3A-D). When focusing the analyses on key enzymes that catalyze irreversible reactions (Fig.3E), we observed that vagotomy increased the expression of proteins involved in glycolysis (Gck, Pfkl, Aldob, Pklr), fatty acid biosynthesis initiation and elongation (Fasn, Acaca, Acacb, Mcat), Fatty acid elongation in endoplasmic reticulum (Scd1 and Acot1), as well as ω-oxidation (Cyp4a14). Strikingly, the expression of key proteins involved in β-oxidation (Acox1 and Acaa1a) and gluconeogenesis (Fbp1 and Pck1) were reduced following VNX. These results point to a major alteration in the hepatic metabolic flux towards glycolysis and fatty acid biosynthesis.

**Figure 3:**
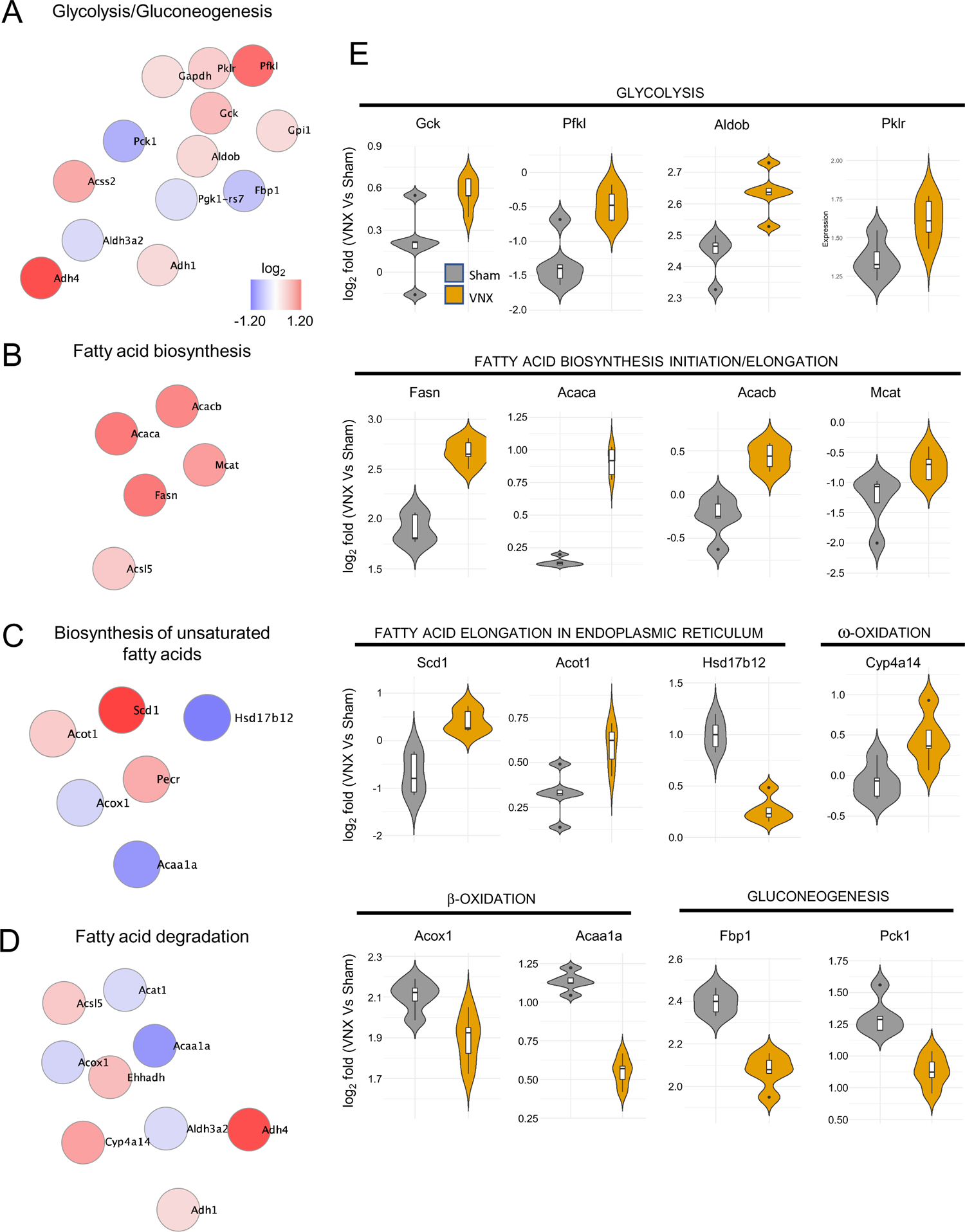
Liver metabolic shift following vagotomy. (**A-D)** Interaction network of differentially abundant proteins (p<0.05, FDR adjusted-p<0.05) within the selected enriched KEGG pathways. **(E)** Violin plots of the relative expression of the proteins grouped by physiological actions.

### 3.4. The alterations in hepatic metabolic machinery are independent of major changes in GI tract physiology

The gastrointestinal tract is an important target of vagus nerve innervation. Therefore, we raised the possibility that the metabolic alterations in the liver shown in the proteomic analysis could be a consequence of changes in gastrointestinal physiology. To investigate this hypothesis, we first monitored the total locomotor activity of sham and VNX animals, using telemetry. We observed that animals of the two groups presented similar locomotor patterns throughout the day (Fig.4A) as well as food consumption before and after the surgery (Fig.4B), pointing to no major changes in feeding behavior. Furthermore, we did not observe significant differences in intestinal motility or intestinal absorption following vagotomy (Fig.4C-D). Our results suggest that the metabolic shift in mice submitted to VNX was not paralleled by major changes in nutrient handling in the gastrointestinal tract.

**Figure 4:**
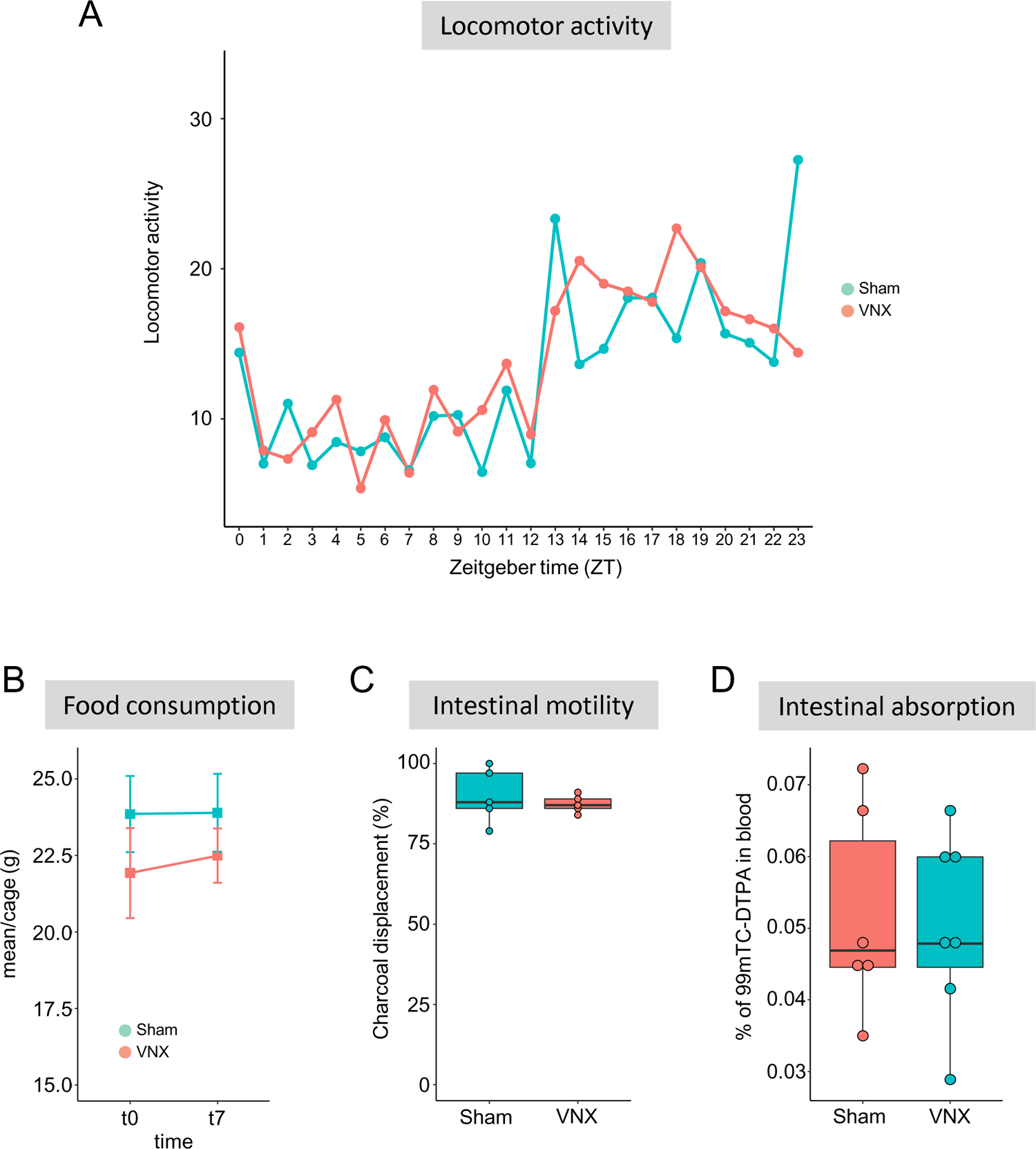
Physiological parameters following vagotomy. **(A)** Locomotor activity, **(B)** Food consumption, **(C)** Intestinal motility and **(D)** Intestinal absorption. VNX = vagotomy; n=5/group in each experiment.

### 3.5. The metabolic shift induced by vagotomy primes the development and severity of NAFLD

Based on our findings that vagotomy shifts liver metabolism towards lipid accumulation by increasing glycolytic and fatty acid biosynthesis fluxes as well as reducing β-oxidation, we asked whether the disruption of vagus nerve circuitry would play a role in the development of NAFLD.

For this purpose, we submitted Sham and VNX mice to a carbohydrate-rich dietary challenge (HC) and addressed lipid accumulation under physiological conditions by using high-definition intravital microscopy (Fig.5A). We found that HC-challenged mice presented a massive accumulation of lipid droplets within the hepatic parenchyma when compared to the group fed with a standard diet (Fig.5B-C). Vagotomy worsened liver lipid accumulation when compared to the Sham group, as observed by the increase of parenchymal area occupied by lipid droplets. Surprisingly, vagotomy resulted in hepatic lipid accumulation even in standard diet-fed animals, corroborating and validating our proteomics data. Food consumption and body weight were not affected by any of the experimental procedures throughout the experiments (Fig.5D-E). We also performed a different dietary challenge using a high-fat diet (HFD) (Fig.6A) and observed similar results. HFD challenge resulted in liver lipid accumulation, which was more severe following VNX (Fig.6B-C). Importantly, the livers of mice submitted to vagotomy and fed a standard diet also displayed more lipid droplets when compared to the livers of the Sham group fed a similar diet. No alterations in body weight were detected (Fig.6D).

**Figure 5:**
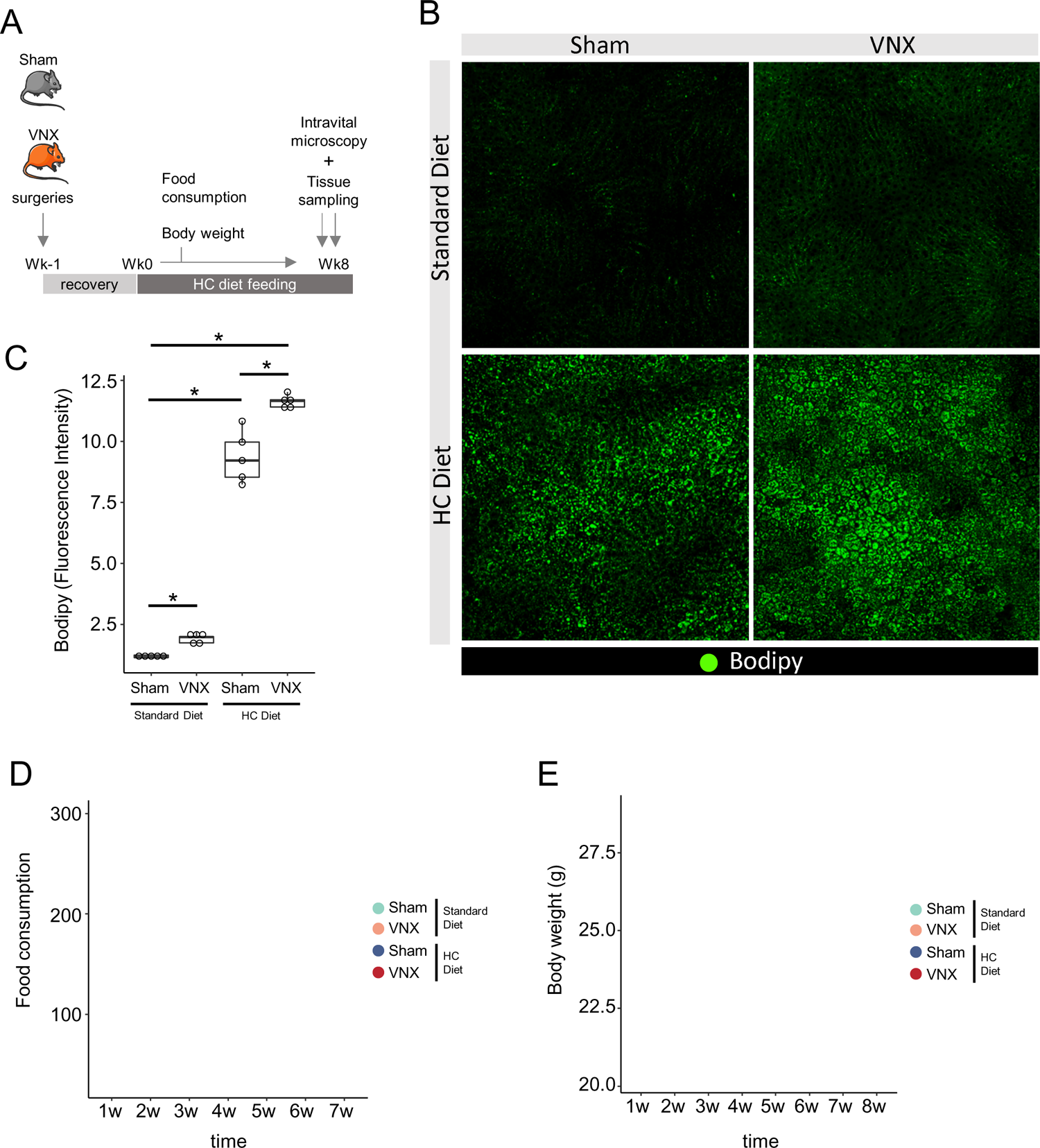
Vagotomy results in increased lipid accumulation in the liver following a high-carbohydrate (HC) dietary challenge. **(A)** Experimental design and protocol. **(B)** Representative *in vivo* confocal images of lipid accumulation assays using Bodipy. **(C)** Quantification of lipids in the liver of Sham and VNX mice (n=5/group, *p<0.05, One Way ANOVA with Newman-Keuls Multiple Comparison Test). **(D-E)** Food consumption (D) and Body weight (E) of mice throughout the experiment. VNX = vagotomy.

**Figure 6:**
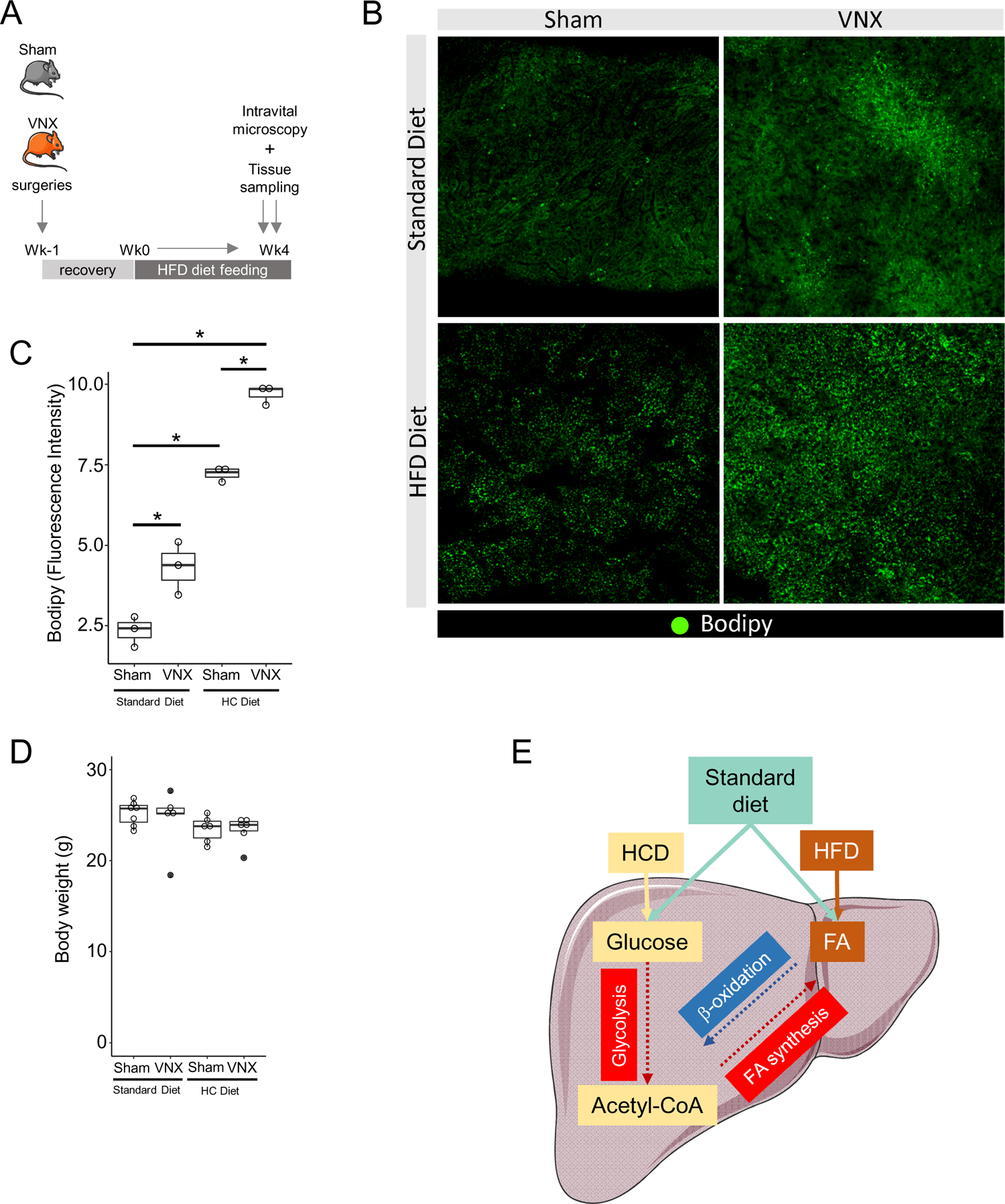
Vagotomy results in increased lipid accumulation in the liver following a high-fat (HFD) dietary challenge. **(A)** Experimental design. **(B)** Representative *in vivo* confocal images of lipid accumulation assays using Bodipy. **(C)** Quantification of lipids in the liver of Sham and VNX mice (n=3/group, *p<0.05, One Way ANOVA with Newman-Keuls Multiple Comparison Test). **(D)** Body weight of mice at the end of experiment. VNX = vagotomy. **(E)** Proposed model. In physiological conditions, the vagus nerve tunes carbohydrate and fatty acid metabolism to maintain metabolic homeostasis. When this circuitry is disrupted, the glycolytic and fatty acid synthesis processes are increased (red rectangles and arrows), whereas β-oxidation is reduced (blue rectangle and arrow). This creates a favorable metabolic flux that primes the liver to accumulate lipids. During a dietary challenge, such as high-carbohydrate diet (HCD) or high-fat diet (HFD), the high nutrient influx in the context of a metabolic shifted liver results in worsened hepatic steatosis.

Taken together, our results show that the metabolic remodeling following vagotomy primes the liver to accumulate lipids, both under a standard diet feeding condition or during different dietary challenges.

## 4. Discussion

The liver is a central organ involved in carbohydrate and lipid metabolism, serving as a major energetic reservoir for other tissues [27] and the regulation of hepatic metabolic homeostasis is pivotal to whole-body metabolism. This balance is partially dependent on the sensing of metabolic states or nutrient availability and the release of neuroendocrine signals [28,29]. Most of these neural circuits are intrinsic to the vagus nerve, with vagal afferent fibers conveying sensory information as mechanical stretching, nutrients/metabolites and hormones, from gastrointestinal tract organs to the nucleus of the solitary tract (NTS) within the central nervous system [7,13,14,30–33]. The responses are then transmitted to the periphery through the efferent fibers of the nerve that originate predominantly from the dorsal motor nucleus of the vagus (DMN) and the ambiguus nucleus (NA) [30].

In this study, we showed that liver metabolic homeostasis is tightly regulated by the vagus nerve, as surgical ablation of this signaling led to a significant alteration of the hepatic proteome landscape. Among the differentially represented pathways, the ones that drew the most attention were those involved in metabolic networks that make up the biochemical processes involved in carbohydrates and lipids metabolism of the liver, such as glycolysis/gluconeogenesis and fatty acid metabolism. Interestingly, the metabolic alterations were not paralleled by changes in food consumption, intestinal transit, or absorption, suggesting that nutrient processing fluxes within the hepatic microenvironment are different following impairment of the vagal circuitry. As a consequence, vagotomized mice showed mild lipid accumulation within hepatocytes even when fed a standard diet. We also observed that steatosis worsened in vagotomized animals following dietary challenges with high-carbohydrate or high-fat diets likely due to a high nutrient influx in the context of a metabolic shifted liver (Fig.5E). The brain-vagus-liver axis was shown to protect the liver against ectopic lipid accumulation by mediating the actions of leptin within the central nervous system, resulting in the activation of the dorsal vagal complex and in increased triglyceride secretion by the liver. [34]. These findings are in line with our main results, showing that the vagus nerve directly shapes liver metabolic function, specially carbohydrate and lipid metabolism. Importantly, this axis also prevented liver steatosis without changes in body weight, food intake, and circulating triglyceride levels [34], corroborating our findings. Recently, the importance of leptin signaling through vagus nerve in preventing liver steatosis was also demonstrated in humans [35].

Vagus nerve neural circuit is a major component of cholinergic signaling, a pathway that has been associated with the control of metabolism. Acetylcholine, the main neurotransmitter released by efferent vagal fibers, exerts its functions interacting with cholinergic receptors, including nicotinic and muscarinic receptors, both of which are expressed in the liver. In line with our findings, mice lacking the nicotinic acetylcholine receptor ⍺7 subunit (⍺7nAChR^-/-^) showed more aggravated hepatic lipid accumulation and steatosis when fed a high-fat diet when compared with WT mice fed with the same diet. Remarkably, the pharmacological activation of ⍺7nAChR in WT mice following a nutritional challenge partially reversed steatosis [36], an effect that involves the regulation of lipogenesis via the regulation of fatty acid synthase (Fasn) expression [37]. Independent studies also showed that impairment of cholinergic signaling mediated by ⍺7nAChR worsened lipid accumulation within the liver in a diet induced model of non-alcoholic steatohepatitis (NASH) [38] or led to metabolic disorders [39]. Our data significantly expands this knowledge, as we show a complex alteration in the hepatic metabolic flux in the context of impaired cholinergic signaling, with the increase in the expression of glycolytic and fatty acid biosynthesis enzymes, concomitant with a decrease in key enzymes of the beta-oxidation pathways (Fig.5E).

Chronic high-calorie ingestion reduces vagal excitability and decreases signaling to the central nervous system [40,41]. Such disruption is sufficient to promote overconsumption of food and weight gain [42,43] and, as described here, likely leads to alteration in the metabolic protein network within the liver. Altogether, these findings suggest vagus nerve-mediated signaling disruption as a putative important event for the onset of metabolic disorders. Within the spectrum of metabolic diseases, non-alcoholic fatty liver disease (NAFLD) is recognized as a major health burden worldwide [44]. NAFLD is a condition characterized by the initial accumulation of fat in the liver that can progress to inflammation and fibrosis (NASH) and that is the fastest-growing indication for liver transplantation [45]. Interestingly, the frequency of *de novo* NAFLD ranges between 18% and 30% in patients submitted to liver transplant [45–47] and, as in the general population, it occurs in the context of altered metabolism, suggesting the major relevance of host factors [48]. Because reinnervation of hepatic parenchyma is limited after transplantation [49], our findings may provide new insights into this condition as well.

## 5. Conclusions

In conclusion, this study describes the liver-brain axis mediated by the vagus nerve as an important regulator of the hepatic metabolic landscape. We found that the disruption of this neural circuit leads to a metabolic shift in the liver towards fatty acid biosynthesis, which primes the organ to accumulate lipids.

## Acknowledgements

This study was funded by INCT-CNPq (#406792/2022-4), CNPq and CAPES and FAPEMIG. The graphical abstract and Figure 5E were partly generated using Servier Medical Art, provided by Servier, licensed under a Creative Commons Attribution 3.0 unported license with minimal changes in opacity, color palette, or text annotation to better suit the message of the text.

Proteomics and mass spectrometry infrastructure at the University of Southern Denmark SDU were supported by generous grants to the VILLUM Center for Bioanalytical Sciences (VILLUM Foundation grant no. 7292), PRO-MS: Danish National Mass Spectrometry Platform for Functional Proteomics (grant no. 5072-00007B), and the Novo Nordisk Foundation (INTEGRA, NNF20OC0061575).

## 6. Author contributions

CFB, RCF, BFV, and GTS conducted all the experiments, and analyzed all the data in this study, with input from other authors. LRR, VG, FK, and TVB designed and performed the proteomics experiments and analysis and data curation, with input from CFB and AGO. ABD, GBM, and CFB conducted the *in vivo* imaging experiments through intravital microscopy. PF, MPO, and CFB performed the surgeries and the locomotor activity experiments. RMP, VNC, SOAF, and CFB performed the absorption experiments with input from AGO. ACCO, AVMF, CFB, and AGO designed the dietary challenges with high-refined carbohydrate or high-fat diets; AVMFL, and JIAL provided the materials and reagents to prepare the diets; ACCO and NAM performed the experiments. AGO conceived the research idea, and supervised the study design as well as all the experiments and data analysis. All authors contributed significantly to the interpretation of data. AGO, CFB, TVB and LRR prepared the submitted version of the manuscript (drafting and revising), with contributions from all authors.

## 7. Role of the funding source

The funding sources had no involvement in the collection, analysis, and interpretation of the data, writing of the manuscript, or in the decision to submit the paper for publication.

## 8. Declaration of interest

Authors declare no competing interests.

## 9. Data availability

The mass spectrometry proteomics data have been deposited to the ProteomeXchange Consortium via the PRIDE [50] partner repository with the dataset identifier PXD041052.

## Abbreviations

ZT: Zeitgeber Time

TEAB: Triethylammonium Bicarbonate

TCEP: Tris(2-carboxyethyl)phosphine

Hz: Hertz

ACN: Acetonitrile

TFA: Trifluoroacetic Acid

VNX: Vagotomy

MS: Mass Spectometry

HCD: High-carbohydrate diet

HFD: High-fat diet

KEGG: Kyoto Encyclopedia of Genes and Genomes

TCA: Tricarboxylic Acid Cycle

DMSO: dimethyl sulfoxide

Gck: Glucokinase

Pfkl: Phosphofructokinase

Aldob: Aldolase, fructose-bisphosphate B

Pklr: Pyruvate kinase liver and red blood cell

Fasn: Fatty acid synthase

Acaca: Acetyl-Coenzyme A carboxylase alpha

Acacb: Acetyl-Coenzyme A carboxylase beta

Mcat: Malonyl CoA:ACP acyltransferase (mitochondrial)

Scd1: Stearoyl-Coenzyme A desaturase 1

Acot1: Acyl-CoA thioesterase 1

Hsd17b12: hydroxysteroid (17-beta) dehydrogenase 12

Cyp4a14: Cytochrome P450, family 4, subfamily a, polypeptide 14

Acox1: Acyl-Coenzyme A oxidase 1, palmitoyl

Acaa1a: Acetyl-Coenzyme A acyltransferase 1A

Fbp1: Fructose bisphosphatase 1

Pck1: Phosphoenolpyruvate carboxykinase 1, cytosolic

NAFLD: Non-Alcoholic Fatty Liver Disease

NASH: Non-Alcoholic Steatohepatitis

NTS: Nucleus of the solitary tract

DMN: Dorsal motor nucleus of the vagus

NA: Ambiguus nucleus

PCA: Principal Component Analysis

FDR: False Discovery Rate

## Notes

### Competing Interest Statement

The authors have declared no competing interest.

## References

[1] Judge A, Dodd MS. Metabolism. Essays Biochem 2020;64:607–47. 10.1042/EBC20190041.

[2] Kietzmann T. Metabolic zonation of the liver: The oxygen gradient revisited. Redox Biol 2017;11:622–30. 10.1016/j.redox.2017.01.012.

[3] Rui L. Energy metabolism in the liver. Compr Physiol 2014;4:177–97. 10.1002/cphy.c130024.

[4] Jensen KJ, Alpini G, Glaser S. Hepatic nervous system and neurobiology of the liver. Compr Physiol 2013;3:655–66. 10.1002/cphy.c120018.

[5] Berthoud H-R. Anatomy and function of sensory hepatic nerves. Anat Rec 2004;280A:827–35. 10.1002/ar.a.20088.

[6] Bohórquez D V., Shahid RA, Erdmann A, Kreger AM, Wang Y, Calakos N, et al. Neuroepithelial circuit formed by innervation of sensory enteroendocrine cells. J Clin Invest 2015;125:782–6. 10.1172/JCI78361.

[7] Kaelberer MM, Buchanan KL, Klein ME, Barth BB, Montoya MM, Shen X, et al. A gut-brain neural circuit for nutrient sensory transduction. Science (80-) 2018;361. 10.1126/science.aat5236.

[8] Niijima A. Glucose-sensitive afferent nerve fibres in the hepatic branch of the vagus nerve in the guinea-pig. J Physiol 1982;332:315–23. 10.1113/jphysiol.1982.sp014415.

[9] Niijima A. Reflex effects of oral, gastrointestinal and hepatoportal glutamate sensors on vagal nerve activity. J. Nutr., vol. 130, 2000. 10.1093/jn/130.4.971s.

[10] Randich A, Spraggins DS, Cox JE, Meller ST, Kelm GR. Jejunal or portal vein infusions of lipids increase hepatic vagal afferent activity. Neuroreport 2001;12:3101–5. 10.1097/00001756-200110080-00024.

[11] Yi CX, la Fleur SE, Fliers E, Kalsbeek A. The role of the autonomic nervous liver innervation in the control of energy metabolism. Biochim Biophys Acta - Mol Basis Dis 2010;1802:416–31. 10.1016/j.bbadis.2010.01.006.

[12] Han W, Tellez LA, Perkins MH, Perez IO, Qu T, Ferreira J, et al. A Neural Circuit for Gut-Induced Reward. Cell 2018;175:665–678.e23. 10.1016/j.cell.2018.08.049.

[13] Buchanan KL, Rupprecht LE, Kaelberer MM, Sahasrabudhe A, Klein ME, Villalobos JA, et al. The preference for sugar over sweetener depends on a gut sensor cell. Nat Neurosci 2022;25:191–200. 10.1038/s41593-021-00982-7.

[14] de Lartigue G. Role of the vagus nerve in the development and treatment of diet-induced obesity. J Physiol 2016;594:5791–815. 10.1113/JP271538.

[15] Burneo JG, Faught E, Knowlton R, Morawetz R, Kuzniecky R. Weight loss associated with vagus nerve stimulation. Neurology 2002;59:463–4. 10.1212/WNL.59.3.463.

[16] Pardo J V., Sheikh SA, Kuskowski MA, Surerus-Johnson C, Hagen MC, Lee JT, et al. Weight loss during chronic, cervical vagus nerve stimulation in depressed patients with obesity: An observation. Int J Obes 2007;31:1756–9. 10.1038/sj.ijo.0803666.

[17] Vijgen GHEJ, Bouvy ND, Leenen L, Rijkers K, Cornips E, Majoie M, et al. Vagus Nerve Stimulation Increases Energy Expenditure: Relation to Brown Adipose Tissue Activity. PLoS One 2013;8. 10.1371/journal.pone.0077221.

[18] Cretenet G, Le Clech M, Gachon F. Circadian Clock-Coordinated 12 Hr Period Rhythmic Activation of the IRE1α Pathway Controls Lipid Metabolism in Mouse Liver. Cell Metab 2010;11:47–57. 10.1016/j.cmet.2009.11.002.

[19] Fonseca RC, Bassi GS, Brito CC, Rosa LB, David BA, Araújo AM, et al. Vagus nerve regulates the phagocytic and secretory activity of resident macrophages in the liver. Brain Behav Immun 2019;81:444–54. 10.1016/j.bbi.2019.06.041.

[20] Boersema PJ, Raijmakers R, Lemeer S, Mohammed S, Heck AJR. Multiplex peptide stable isotope dimethyl labeling for quantitative proteomics. Nat Protoc 2009;4:484–94. 10.1038/nprot.2009.21.

[21] Cox J, Neuhauser N, Michalski A, Scheltema RA, Olsen J V., Mann M. Andromeda: A peptide search engine integrated into the MaxQuant environment. J Proteome Res 2011;10:1794–805. 10.1021/pr101065j.

[22] Darzi Y, Yamate Y, Yamada T. FuncTree2: An interactive radial tree for functional hierarchies and omics data visualization. Bioinformatics 2019;35:4519–21. 10.1093/bioinformatics/btz245.

[23] Yu G, Wang LG, Han Y, He QY. ClusterProfiler: An R package for comparing biological themes among gene clusters. Omi A J Integr Biol 2012;16:284–7. 10.1089/omi.2011.0118.

[24] Bindea G, Mlecnik B, Hackl H, Charoentong P, Tosolini M, Kirilovsky A, et al. ClueGO: A Cytoscape plug-in to decipher functionally grouped gene ontology and pathway annotation networks. Bioinformatics 2009;25:1091–3. 10.1093/bioinformatics/btp101.

[25] Szklarczyk D, Kirsch R, Koutrouli M, Nastou K, Mehryary F, Hachilif R, et al. The STRING database in 2023: protein-protein association networks and functional enrichment analyses for any sequenced genome of interest. Nucleic Acids Res 2023;51:D638–46. 10.1093/nar/gkac1000.

[26] David BA, Rezende RM, Antunes MM, Santos MM, Freitas Lopes MA, Diniz AB, et al. Combination of Mass Cytometry and Imaging Analysis Reveals Origin, Location, and Functional Repopulation of Liver Myeloid Cells in Mice. Gastroenterology 2016;151:1176–91. 10.1053/j.gastro.2016.08.024.

[27] Gardemann A, Püschel Gp, Jungermann K. Nervous control of liver metabolism and hemodynamics. Eur J Biochem 1992;207:399–411. 10.1111/j.1432-1033.1992.tb17063.x.

[28] Seicol BJ, Bejarano S, Behnke N, Guo L. Neuromodulation of metabolic functions: From pharmaceuticals to bioelectronics to biocircuits. J Biol Eng 2019;13:67. 10.1186/s13036-019-0194-z.

[29] Cabot L, Erlenbeck-Dinkelmann J, Fenselau H. Neural gut-to-brain communication for postprandial control of satiation and glucose metabolism. J Endocrinol 2023;258. 10.1530/JOE-22-0320.

[30] Neuhuber WL, Berthoud HR. Functional anatomy of the vagus system – Emphasis on the somato-visceral interface. Auton Neurosci Basic Clin 2021;236. 10.1016/j.autneu.2021.102887.

[31] Masi EB, Valdés-Ferrer SI, Steinberg BE. The vagus neurometabolic interface and clinical disease. Int J Obes 2018;42:1101–11. 10.1038/s41366-018-0086-1.

[32] Vana V, Lærke MK, Rehfeld JF, Arnold M, Dmytriyeva O, Langhans W, et al. Vagal afferent cholecystokinin receptor activation is required for glucagon-like peptide-1– induced satiation. Diabetes, Obes Metab 2022;24:268–80. 10.1111/dom.14575.

[33] Cook TM, Gavini CK, Jesse J, Aubert G, Gornick E, Bonomo R, et al. Vagal neuron expression of the microbiota-derived metabolite receptor, free fatty acid receptor (FFAR3), is necessary for normal feeding behavior. Mol Metab 2021;54. 10.1016/j.molmet.2021.101350.

[34] Hackl MT, Fürnsinn C, Schuh CM, Krssak M, Carli F, Guerra S, et al. Brain leptin reduces liver lipids by increasing hepatic triglyceride secretion and lowering lipogenesis. Nat Commun 2019;10:2717. 10.1038/s41467-019-10684-1.

[35] Metz M, Beghini M, Wolf P, Pfleger L, Hackl M, Bastian M, et al. Leptin increases hepatic triglyceride export via a vagal mechanism in humans. Cell Metab 2022;34:1719–1731.e5. 10.1016/j.cmet.2022.09.020.

[36] Li DJ, Liu J, Hua X, Fu H, Huang F, Fei YB, et al. Nicotinic acetylcholine receptor α7 subunit improves energy homeostasis and inhibits inflammation in nonalcoholic fatty liver disease. Metabolism 2018;79:52–63. 10.1016/j.metabol.2017.11.002.

[37] Hasan MK, Friedman TC, Sims C, Lee DL, Espinoza-Derout J, Ume A, et al. a7-nicotinic acetylcholine receptor agonist ameliorates nicotine plus high-fat Diet–Induced hepatic steatosis in male mice by inhibiting oxidative stress and stimulating AMPK signaling. Endocrinology 2018;159:931–44. 10.1210/en.2017-00594.

[38] Kimura K, Inaba Y, Watanabe H, Matsukawa T, Matsumoto M, Inoue H. Nicotinic alpha-7 acetylcholine receptor deficiency exacerbates hepatic inflammation and fibrosis in a mouse model of non-alcoholic steatohepatitis. J Diabetes Investig 2019;10:659–66. 10.1111/jdi.12964.

[39] Gausserès B, Liu J, Foppen E, Tourrel-Cuzin C, Sanchez-Archidona AR, Delangre E, et al. The constitutive lack of α7 nicotinic receptor leads to metabolic disorders in mouse. Biomolecules 2020;10:1–28. 10.3390/biom10071057.

[40] Al Helaili A, Park SJ, Beyak MJ. Chronic high fat diet impairs glucagon like peptide-1 sensitivity in vagal afferents. Biochem Biophys Res Commun 2020;533:110–7. 10.1016/j.bbrc.2020.08.045.

[41] Loper H, Leinen M, Bassoff L, Sample J, Romero-Ortega M, Gustafson KJ, et al. Both high fat and high carbohydrate diets impair vagus nerve signaling of satiety. Sci Rep 2021;11. 10.1038/s41598-021-89465-0.

[42] de Lartigue G, Ronveaux CC, Raybould HE. Deletion of leptin signaling in vagal afferent neurons results in hyperphagia and obesity. Mol Metab 2014;3:595–607. 10.1016/j.molmet.2014.06.003.

[43] Barella LF, Miranda RA, Franco CCS, Alves VS, Malta A, Ribeiro TAS, et al. Vagus nerve contributes to metabolic syndrome in high-fat diet-fed young and adult rats. Exp Physiol 2015;100:57–68. 10.1113/EXPPHYSIOL.2014.082982.

[44] Yki-Järvinen H. Non-alcoholic fatty liver disease as a cause and a consequence of metabolic syndrome. Lancet Diabetes Endocrinol 2014;2:901–10. 10.1016/S2213-8587(14)70032-4.

[45] Germani G, Laryea M, Rubbia-Brandt L, Egawa H, Burra P, O’Grady J, et al. Management of Recurrent and de Novo NAFLD/NASH after Liver Transplantation. Transplantation 2019;103:57–67. 10.1097/TP.0000000000002485.

[46] Seo S, Maganti K, Khehra M, Ramsamooj R, Tsodikov A, Bowlus C, et al. De novo nonalcoholic fatty liver disease after liver transplantation. Liver Transplant 2007;13:844–7. 10.1002/lt.20932.

[47] Dumortier J, Giostra E, Belbouab S, Morard I, Guillaud O, Spahr L, et al. Non-alcoholic fatty liver disease in liver transplant recipients: Another story of seed and soil. Am J Gastroenterol 2010;105:613–20. 10.1038/ajg.2009.717.

[48] Gitto S, De Maria N, Di Benedetto F, Tarantino G, Serra V, Maroni L, et al. De-novo nonalcoholic steatohepatitis is associated with long-term increased mortality in liver transplant recipients. Eur J Gastroenterol Hepatol 2018;30:766–73. 10.1097/MEG.0000000000001105.

[49] Boon AP, Hubscher SG, Lee JA, Hines JE, Burt AD. Hepatic reinnervation following orthotopic liver transplantation in man. J Pathol 1992;167:217–22. 10.1002/path.1711670210.

[50] Perez-Riverol Y, Bai J, Bandla C, García-Seisdedos D, Hewapathirana S, Kamatchinathan S, et al. The PRIDE database resources in 2022: A hub for mass spectrometry-based proteomics evidences. Nucleic Acids Res 2022;50:D543–52. 10.1093/nar/gkab1038.

